# Single cell transcriptome profiling of the human alcohol-dependent brain

**DOI:** 10.1101/780304

**Authors:** Eric Brenner, Gayatri R. Tiwari, Yunlong Liu, Amy Brock, R. Dayne Mayfield

## Abstract

**Background:** Alcoholism remains a prevalent health concern throughout the world. Previous studies have identified transcriptomic patterns in the brain associated with alcohol dependence in both humans and animal models. But none of these studies have systematically investigated expression within the unique cell types present in the brain.

**Results:** We utilized single nucleus RNA sequencing (snRNA-seq) to examine the transcriptomes of over 16,000 nuclei isolated from prefrontal cortex of alcoholic and control individuals. Each nucleus was assigned to one of seven major cell types by unsupervised clustering. Cell type enrichment patterns varied greatly among neuroinflammatory-related genes, which are known to play roles in alcohol dependence and neurodegeneration. Differential expression analysis identified cell type-specific genes with altered expression in alcoholics. The largest number of differentially expressed genes (DEGs), including both protein-coding and non-coding, were detected in astrocytes, oligodendrocytes, and microglia.

**Conclusions:** To our knowledge, this is the first single cell transcriptome analysis of alcohol-associated gene expression in any species, and the first such analysis in humans for any addictive substance. These findings greatly advance understanding of transcriptomic changes in the brain of alcohol-dependent individuals.

## Background

Alcohol abuse is involved in over 200 pathologies and health conditions (e.g. alcohol dependence, liver cirrhosis, cancers, and injuries) and creates substantial social and economic burdens (1,2). To develop more effective therapeutic strategies, we must first understand how alcohol affects the body at the cellular and molecular level. Previous studies of transcriptomic responses in human alcoholics (3–6) relied on RNA extracted from brain regions using tissue homogenates comprised of multiple cell types. This approach likely masks differences in gene expression patterns among specific cells, as well as heterogeneity from cell-to-cell variation within a given cell type.

Single cell RNA sequencing (scRNA-seq) has recently gained attention in cell and molecular biology research for its ability to profile novel cell types and measure cell-to-cell variation in gene expression. To our knowledge, transcriptomic responses to chronic alcohol exposure or any other abused drug have not been studied at the cellular level in the human brain. We hypothesized that cell type-specific gene expression patterns associated with alcoholism will identify novel alcohol targets that were previously missed by bulk analysis of tissue homogenates, as has been shown for other neuropathologies (7,8). Using single nucleus RNA-seq (snRNA-seq), a popular scRNA-seq alternative for analyzing frozen brain tissue (9–18), we profiled the transcriptomes of 16,305 nuclei extracted from frozen prefrontal cortex (PFC) samples of 4 control and 3 alcohol dependent individuals. The PFC is involved in executive function and is an important substrate in the reward circuitry associated with development of alcohol dependence (19). The PFC has also been the focus of many transcriptomic studies (3,4,20,21) and was a logical choice for our initial work. Using the approach presented here, we discovered novel cell type-specific transcriptome changes associated with alcohol abuse in the human PFC.

## Results and Discussion

Like most organs in the body, the brain consists of a diverse array of cell types. In order to systematically assess the roles of different cells in alcohol dependence, we examined gene expression in the PFC of alcohol dependent versus control donors at the single cell level (**Table S1**). For this specific approach, we utilized droplet-based snRNA-seq technology to prepare libraries from postmortem tissue samples. The dataset was comprised of transcriptomes of 16,305 nuclei from seven donors (four control donors and three donors with alcohol dependence) (**Table S2**). We visualized the data using uniform manifold approximation and projection (UMAP) (22). Nuclei did not separate by the donor or batch from which they were isolated, indicating that any sample-specific gene expression patterns that may be present were of minimal impact (**Fig 1A, Fig S1**).

**Figure 1.**
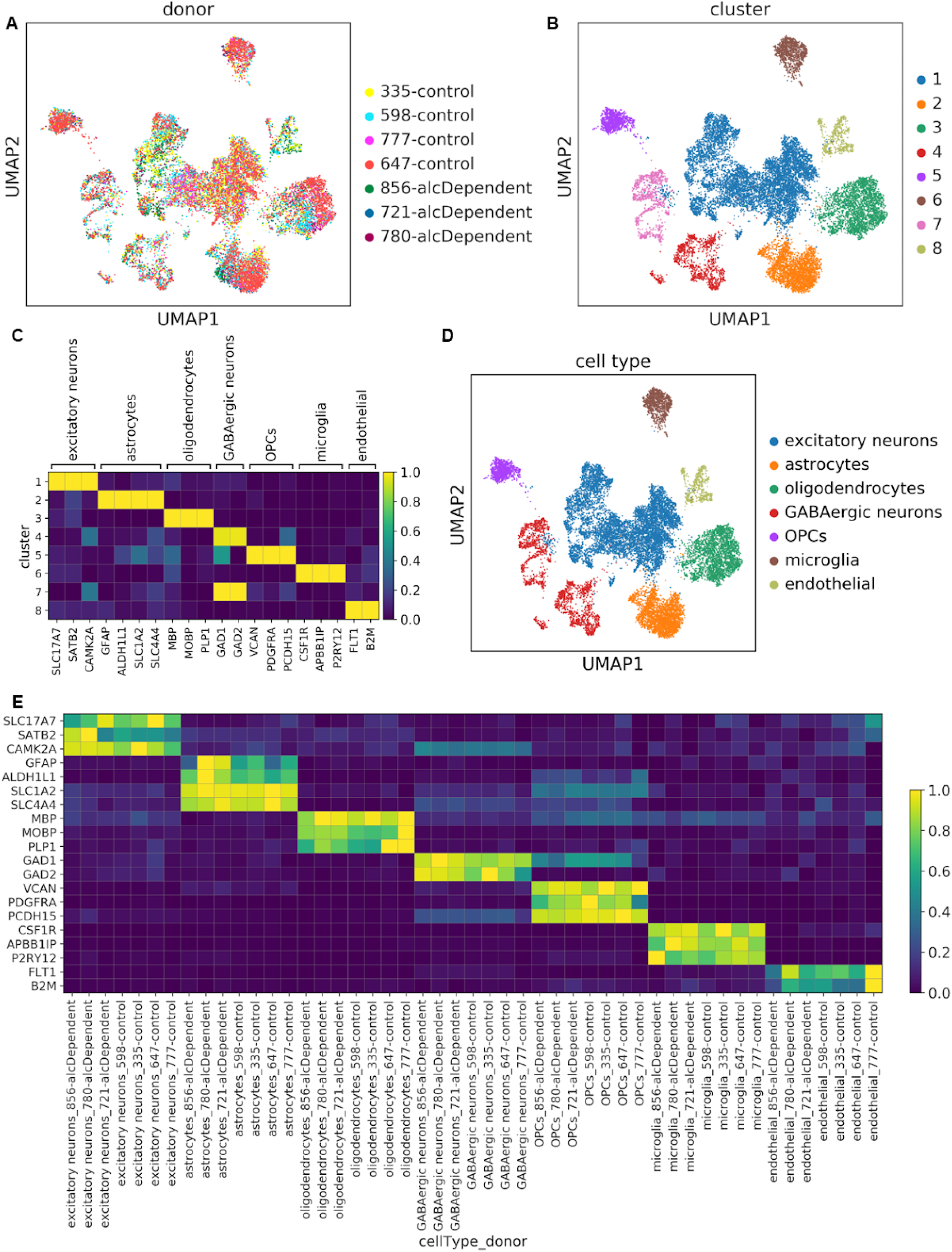
Unsupervised clustering captured expected cell types in every sample. **(A)** UMAP plots of the 16,305 nuclei in our dataset, colored by donor or **(B)** transcriptomically-distinct clusters determined from unsupervised clustering. **(C)** Scaled mean expression of known cell type markers in the different clusters. For each gene, expression was scaled from 0.0-1.0 to maintain a balanced colormap. **(D)** UMAP plot of nuclei colored by cell type assignment. **(E)** Scaled mean expression of marker genes for each cell type in each donor.

### Unsupervised clustering of nuclei

In order to identify transcriptomically-distinct groups of nuclei, we performed unsupervised, graph-based clustering, yielding eight clusters (**Fig 1B**). We then annotated the clusters using the following markers of major brain cell types that are consistently detected in single cell/nucleus transcriptomics studies (7,8,20,23,24): excitatory neurons, astrocytes, oligodendrocytes, inhibitory/GABAergic neurons, oligodendrocyte progenitor cells (OPCs), microglia, and endothelial cells (**Fig 1C-D**). Two clusters corresponded with GABAergic neurons while all other clusters corresponded to a single cell type. We confirmed that the cell types and signatures were conserved among samples from all of the donors by examining marker expression in each donor cell-type combination (**Fig 1E, Table S3**). Proportions of the different cell types were comparable to other snRNA-seq studies (8,23) with neurons (particularly excitatory neurons) being over-represented relative to non-neuronal cell types (**Fig S2**). Furthermore, we did not find differences in cell type proportions between alcoholics and controls (**Fig S2**). This is consistent with previous findings (3).

While some other single cell transcriptomics studies have attempted to define novel cell states and subtypes, our goal was to evaluate gene expression differences between alcoholics and controls in known, established cell types. Therefore, we performed the clustering using a low-resolution parameter (**Methods**) rather than using subclustering.

### Neuroinflammatory signaling

The neuroimmune system is critical for the pathogenesis of neurological diseases such as Alzheimer’s disease and multiple sclerosis (25), and is also involved in regulating alcohol abuse and dependence (26). To date, however, single cell/nucleus RNA-seq studies have not specifically evaluated neuroimmune gene expression among cell types. Of the 85 human genes directly listed in the Neuroinflammation Signaling Pathway in the IPA database, 79 were detected in our snRNA-seq dataset, and 68 were detected in more than 20 nuclei. The expression of these genes among the different cell types is displayed in **Fig 2A**. Expression of some genes was fairly pervasive across cell types (e.g. *HMGB1*) while other genes exhibited relative enrichment in one or two specific cell types (e.g. *SNCA* in excitatory neurons, *BCL2* in astrocytes, *P2RX7* in oligodendrocytes, *GRIA1* in GABAergic neurons, *S100B* in OPCs, *TLR2* in microglia (**Fig 2A-C**), and *CTNNB1* in endothelial cells). Nearly half of the genes whose expression was highest in astrocytes are involved with interferon signaling: *BCL2* (27), *IRF3, HMGB1, TICAM1*, and *TRAF3* (28)

**Figure 2.**
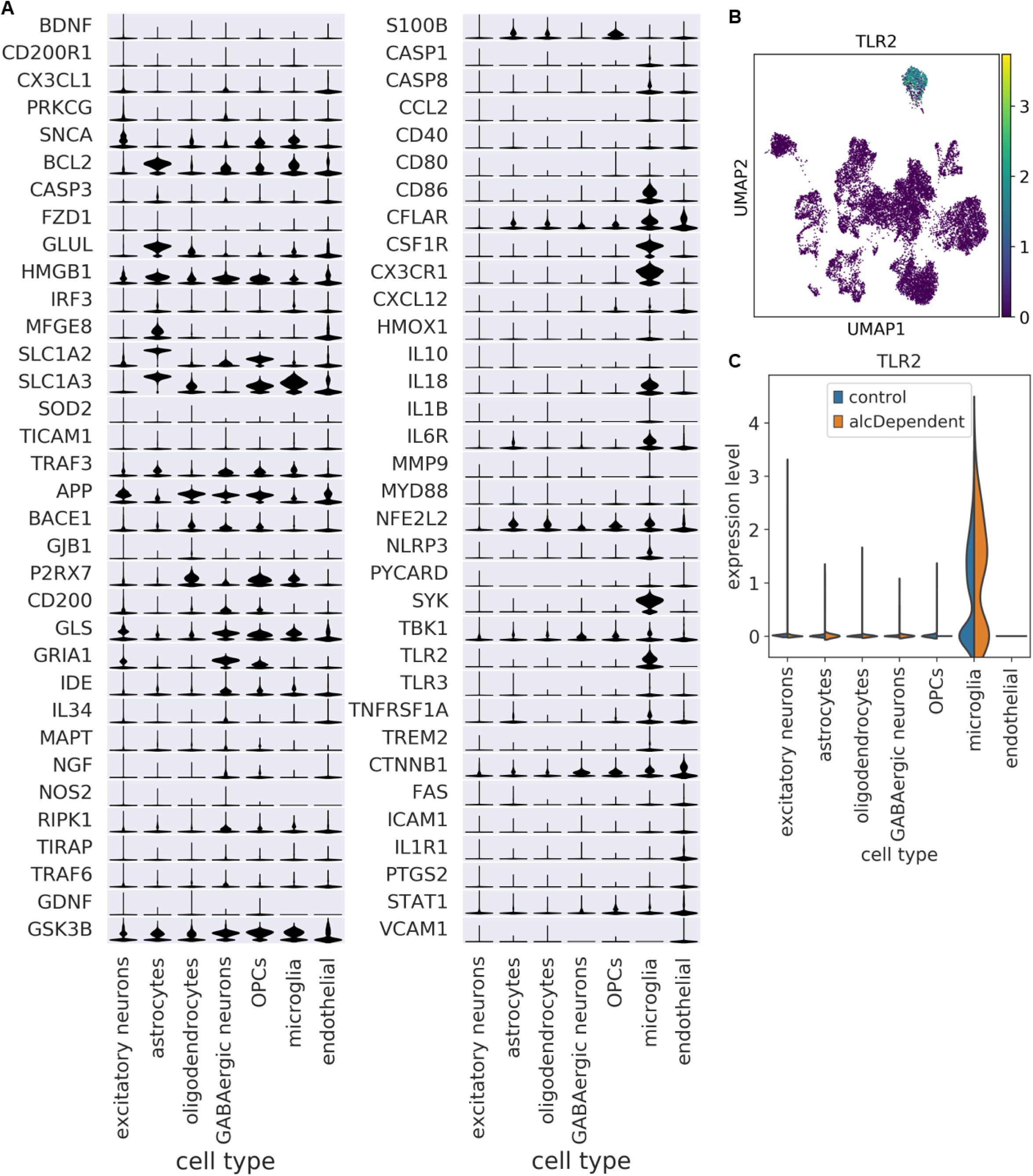
Cell type enrichment patterns vary among neuroimmune genes. **(A)** Scaled expression of neuroimmune genes among cell types. Genes are ordered by which cell type had the highest mean expression. Scaling was done for each gene across all cells. **(B)** UMAP plot depicting expression of TLR2 as an example of a gene that is enriched in a specific cell type (microglia). **(C)** TLR2 expression is enriched in microglia among nuclei from both controls and alcoholics.

While mean expression of a given gene may be higher in a specific cell type, that does not imply that expression in other cell types is statistically negligible or biologically unimportant. For example, expression of *FAS* is lower in astrocytes relative to endothelial cells; however, *FAS* is known to be active in both astrocytes(29) and endothelial cells (30).

### Differential expression and pathway analysis

Prior to differential expression (DE) analysis, we pooled transcript counts within each cell type and each donor, creating “pseudo-bulk” transcriptomes. This approach has been employed by several single cell transcriptomics studies (31–34) since it provides robustness to varying numbers and library sizes of cells/nuclei among replicates (donors acting as replicates in our case), provides strong type I error control, and allows the use of bulk RNA-seq DE tools that have been optimized and validated over the course of many years. After pseudobulking, a typically DESeq2 was used to test for differentially expressed genes (DEGs), with batch as a covariate. We identified a total of 916 DEGs at FDR<0.25 and 253 at FDR<0.05 (**Fig 3A**). There were large differences in DEG counts among the different cell types, with endothelial cells having 0 and astrocytes having 206 DEGs. Recent mouse RNA-seq studies on isolated glial cell types also detected more alcohol-related DEGs in astrocytes than in total brain homogenate (20) and microglia (21). These findings support key roles for astrocytes in response to chronic alcohol that were not detected in other studies lacking cell type resolution.

**Figure 3.**
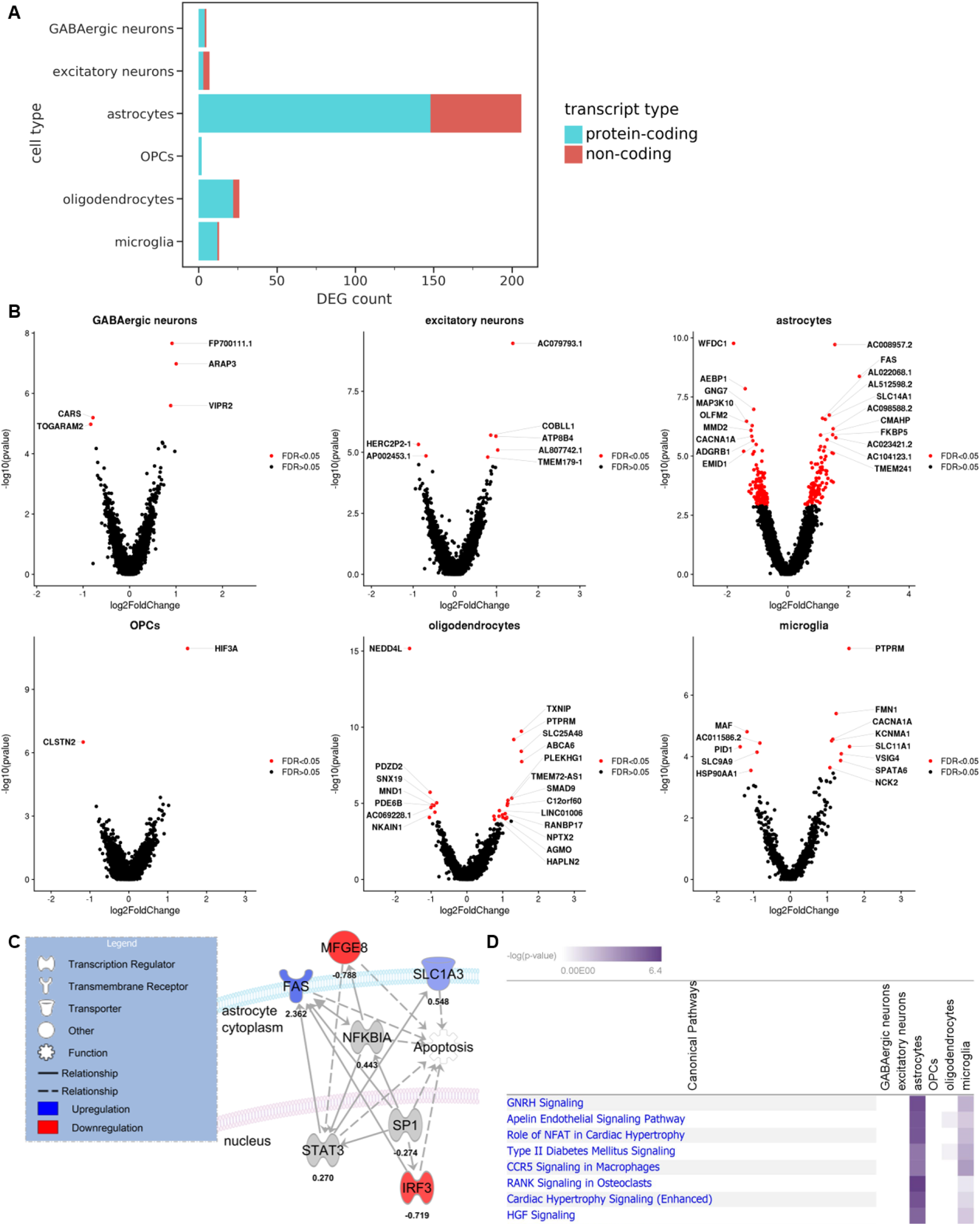
Differentially expressed genes (DEGs) associated with alcoholism were detected in every neural cell type. **(A)** Breakdown of DEGs by cell type and transcript type (FDR<0.05). **(B)** Volcano plots of top DEGs in each cell type (FDR<0.05). **(C)** An Ingenuity Pathway Analysis (IPA) molecular network including four neuroinflammation-associated DEGs in astrocytes (FDR<0.11). Solid lines indicate direct relationships, while dashed lines indicate indirect relationships. **(D)** Log2 fold changes and adjusted p-values of genes were processed by IPA with FDR<0.25 as the cutoff for indicating significant DEGs. The top canonical pathways are shown. Negative log(p-values) are derived from Fisher’s exact test.

DEGs corresponding to both protein-coding and non-coding transcripts were present in every cell type that had more than two total DEGs (**Fig 3A-B**). Bulk and scRNA-seq studies often exclude non-coding RNAs from their analyses (3,8,35). However, the importance of ncRNAs is becoming increasingly apparent, not only in alcohol dependence (36), but in brain physiology in general (37), and these RNAs were therefore included in our data. Interestingly, the top DEG in astrocytes was *AC008957.2*, a lncRNA that is antisense to the gene encoding *SLC1A3*, a glutamate uptake transporter that plays important roles in the neurocircuitry of addiction (38–40). *SLC1A3* was also upregulated in astrocytes, albeit at a lower significance level (FDR=0.07).

In addition to non-coding genes, protein-coding genes of interest were also identified. For example, four DEGs involved in neuroinflammation (*SLC1A3, FAS, MFGE8, IRF3*) were found in astrocytes (**Fig 3C**) (FDR<0.11). IPA revealed that all four are associated with apoptosis, a component of the neuroimmune response. *SLC1A3* (also called “*GLAST*”) was downregulated with alcohol consumption in mouse astrocytes (20,21) but upregulated in the human alcoholic brain (39). *FAS* and *MFGE8*, however, showed similar regulation between mouse and human astrocytes (21).

We also used IPA to evaluate DEGs (FDR<0.25) in all cell types. Only astrocytes, microglia, and oligodendrocytes showed significant pathways (p<0.05, Fisher’s exact test) containing multiple DEGs. The top overall pathway was *GNRH* signaling (**Fig 3D**). One of the top DEGs in this pathway was *CACNA1A*, which was downregulated in astrocytes and upregulated in microglia.

A notable limitation of our DE analysis was the sample size of our dataset (7 donors). A small sample size in a single cell genomics study does affect statistical power, but does not preclude the possibility of drawing meaningful inferences of cell type gene expression in the human brain (9,41). That said, our goal is to increase the sample size for future single cell studies to better identify important expression patterns, particularly for genes that have smaller effect sizes.

### Comparison with previous bulk RNA-seq datasets

We next determined which cell types were most similar or dissimilar to bulk data in terms of alcohol-associated expression changes. We compared our DE results to that from a published bulk RNA-seq analysis of the PFC from 65 alcoholics and 73 control donors (3). Hierarchical clustering revealed that all non-microglia cell types were more similar to bulk than they were to microglia in terms of DE patterns between alcoholics and controls (**Fig 4A**). Other studies have also found that microglial expression, in general, is particularly underrepresented in bulk transcriptomes, relative to that of other brain cell types (8,42). Among the 33 DEGs in microglia, only the gene with the highest log fold change, SLC11A1, had been detected as a significant hit in the bulk data (**Fig 4B**). This suggests that microglia-specific DEGs must have a particularly large effect size in order to be detected in bulk DE analysis

**Figure 4.**
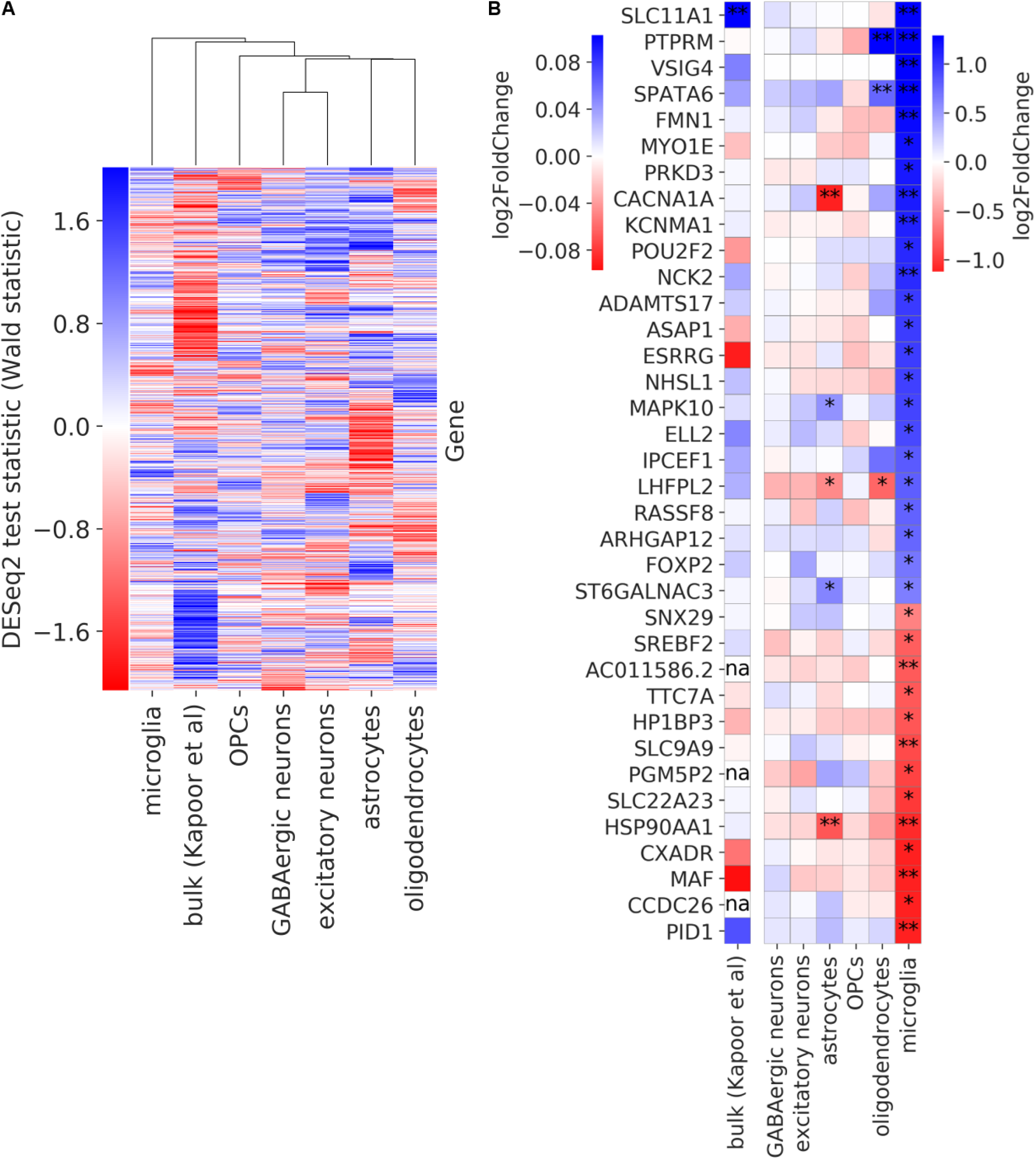
Comparison of alcohol dependence-associated differential expression from human bulk RNA-seq and snRNA-seq data. **(A)** Gene expression changes in alcoholics for each cell type as well as bulk data from a previous study (3). Groups were hierarchically clustered using average as the linkage method. **(B)**. Log fold change of expression in alcohol dependent donors compared with controls for genes that are differentially expressed (FDR<0.25) in microglia. A single asterisk indicates differential expression at FDR<0.25, while a double asterisk indicates FDR<0.05. The three genes with “na” are non-protein-coding genes, which had not been included in the bulk study.

## Conclusions

This study presents the first single cell transcriptomic dataset of the human alcoholic brain. Among 16,305 nuclear transcriptomes, we detected major neural cell types from seven donors: three alcoholics and four controls. Every cell type displayed relative enrichment of different genes linked to neuroinflammation, a process associated with excessive alcohol use. We detected DEGs between alcoholics and controls in every neural cell type. Many more DEGs were found in astrocytes, oligodendrocytes, and microglia relative to neurons, indicating that glial cells should be given particular attention in future studies of alcohol-associated gene expression.

## Methods

### Case selection and postmortem tissue collection

Diagnosis of alcohol dependence was based on DSM-IV criteria. All donors were required to meet the following criteria to be considered in this study: no head injury at time of death, lack of developmental disorder, no recent cerebral stroke, no history of other psychiatric or neurological disorders, and no history of intravenous or polydrug abuse. Alcoholic and control donors were selected to be as close as possible in terms of age, sex, post-mortem interval (PMI), pH of tissue, and cause of death (**Table S1**). Fresh frozen tissue samples were collected as previously described (3).

### Isolation of nuclei from frozen postmortem brain tissue

Nuclei were isolated from tissue based on a modified version of Luciano Martelotto’s Customer-Developed Protocol provided on the 10XGenomics website (43). All procedures were carried out on ice and all centrifugation steps were at 500g for 5 min at 4°C. PFC tissue (∼50 mg) was homogenized in 5 mL of Lysis Buffer [Nuclei EZ Lysis Buffer (Sigma, NUC101), 0.2 U/µL RNase inhibitor (NEB, M0314), 1x protease inhibitor (Sigma, 4693132001)] in 7 mL dounce ∼30 times until no visible tissue pieces remained. An aliquot of homogenate (500 µL) was snap-frozen on dry ice and stored at -80 °C for later RNA integrity assessment. The remaining homogenate was incubated on ice for 5-10 minutes and then centrifuged. The supernatant was removed, and nuclei were resuspended in 2 mL of Lysis Buffer without protease inhibitor. The suspension was incubated for another 5 min on ice and centrifuged. Supernatant was removed, and nuclei were resuspended in 2 mL of Wash & Resuspension Buffer (PBS with 1% BSA and 0.2 U/µL of RNase inhibitor). Nuclei were centrifuged and the supernatant was removed. Nuclei were then resuspended in Wash & Resuspension Buffer containing 10 µg/mL of DAPI and filtered into a FACS tube with a 35-µm strainer cap. Nuclei were then sorted into a tube containing 10XGenomics (v3.0) reverse transcription (RT) master mix (20 µL of RT Reagent, 3.1 µL of Template Switch Oligo, 2 µL of Reducing Agent B, and 10 µL of water). Nuclei-RT mix was topped off to 71.7 µL using water, and then 8.3 µL of RT Enzyme C was added. It should be noted that the sample 647 was prepared in another batch using a slightly different protocol, the main difference being that nuclei were sorted into an empty tube, concentrated by centrifuging and removing all but 100 µL of supernatant, and then resuspended before being combined with the RT master mix.

### RNA integrity assessment

RNA was extracted from the homogenates with a RNeasy Lipid Tissue Mini Kit (Qiagen,74804), using the manufacturer’s instructions. The RNA was quantified using a NanoDrop1000 (Thermo Fisher) and assayed for quality using an Agilent 2100 TapeStation (Agilent Technologies). Homogenate for sample 647 had not been saved.

### Droplet-based snRNA-seq

Nuclei-RT mix (75 µL) was transferred to a Chromium B Chip, and libraries were prepared using the 10XGenomics 3’ Single Cell Gene Expression protocol. Paired-end sequencing of the libraries was conducted using a NovaSeq 6000 with an S2 chip (100 cycles).

### Alignment to reference genome

A GENCODE v30 GTF file was modified to create a “pre-mRNA” GTF file so that pre-mRNAs would be included as counts in the subsequent analysis. Cellranger’s (v3.0.2) *mkref* command was then used to create a pre-mRNA reference from the GTF file and a FASTA file of the GRCh38.p12 genome. FASTQ files of the snRNA-seq libraries were then aligned to the pre-mRNA reference using the *cellranger count* command, producing gene expression matrices. The matrices for the different samples were concatenated into a single matrix using the *cellranger aggr* command with normalization turned off, so that the raw counts would remain unchanged at this point.

### Filtering and normalization

The filtered matrix produced by cellranger was loaded into scanpy (v1.4.3) (44). Nuclei with more than 20% of UMI (transcript) counts attributed to mitochondrial genes were removed, and then all mitochondrial genes were removed from the dataset. Normalization was conducted based on the recommendations from multiple studies that compared several normalization techniques (35,45,46). In brief, three steps were performed: (1) preliminary clustering of cells by constructing a nearest network graph and using scanpy’s implementation of Louvain community detection, (2) calculating size factors using the R package scran (v1.10.2) (47), and (3) dividing counts by the respective size factor assigned to each cell. Normalized counts were then transformed by adding a pseudocount of 1 and taking the natural log.

### Assigning nuclei to transcriptomically-distinct clusters

Genes that are highly variable within every sample were identified using scanpy’s *highly_variable_genes* function with sample identity used for the *batch_key* argument. This type of approach helps select genes that distinguish cell types from each other (9). Normalized expression of every gene was then centered to a mean of zero. Principal component (PC) analysis was performed on the highly variable genes, and the top 50 PCs were used to construct a nearest neighbor graph. Nuclei were then assigned to clusters using the Louvain algorithm with the resolution set to 0.1.

### Differential gene expression analysis

DE testing was performed separately on each cell type. We adapted a transcript count summation strategy (31) (also called “pseudobulking”). First, nuclei corresponding to the given cell type were selected from the full dataset. Second, raw counts were summed in order to produce a “pseudobulk” transcriptome for each donor. Third, DEGs between the alcohol-dependent and control donors were detected using DESeq2 (v1.24.0) (48) with batch as a covariate.

### Pathway analyses of differential expression

Qiagen’s Ingenuity Pathway Analysis (IPA) software was used to identify canonical pathways associated with DEGs in each cell type. A cutoff of FDR<0.25 was used to separate DEGs from non-DEGs.

## Declarations

### Ethics approval and consent to participate

Not applicable

### Availability of data and materials

Raw and processed data will be made publicly available on the Gene Expression Omnibus (GEO) by the time of publication.

### Competing interests

We have no competing interests to declare for this study.

### Funding

This study was supported by NIH grants U01-AA020926 (RDM) and R01-AA012404 (RDM). EB receives support from two internal fellowships at the University of Texas at Austin: the Provost’s Graduate Excellence Fellowship and the F.M. Jones & H.L. Bruce Endowed Graduate Fellowship in Addiction Science.

### Authors’ contributions

This study was designed by EB, AB, and RDM. RDM and GRT procured and prepared the tissue. EB performed the nuclei isolation with assistance from GRT. YL coordinated the sequencing of the final cDNA libraries. EB performed the data analysis with input from GRT, AB, and RDM. EB wrote the manuscript with editing by GRT, AB, and RDM. All authors read and approved the final manuscript.

## Acknowledgments

The authors thank The University of Texas at Austin’s Genomic Sequencing and Analysis Facility (GSAF) and its members for preparing the cDNA libraries and the Center for Medical Genomics at Indiana University School of Medicine for sequencing the libraries. We are also grateful to the New South Wales Tissue Resource Center at the University of Sydney for providing human brain samples; the Centre is supported by the National Health and Medical Research Council of Australia, Schizophrenia Research Institute, and National Institute on Alcohol Abuse and Alcoholism (NIH/NIAAA R24AA012725).

---

**Supplementary figure 1.**
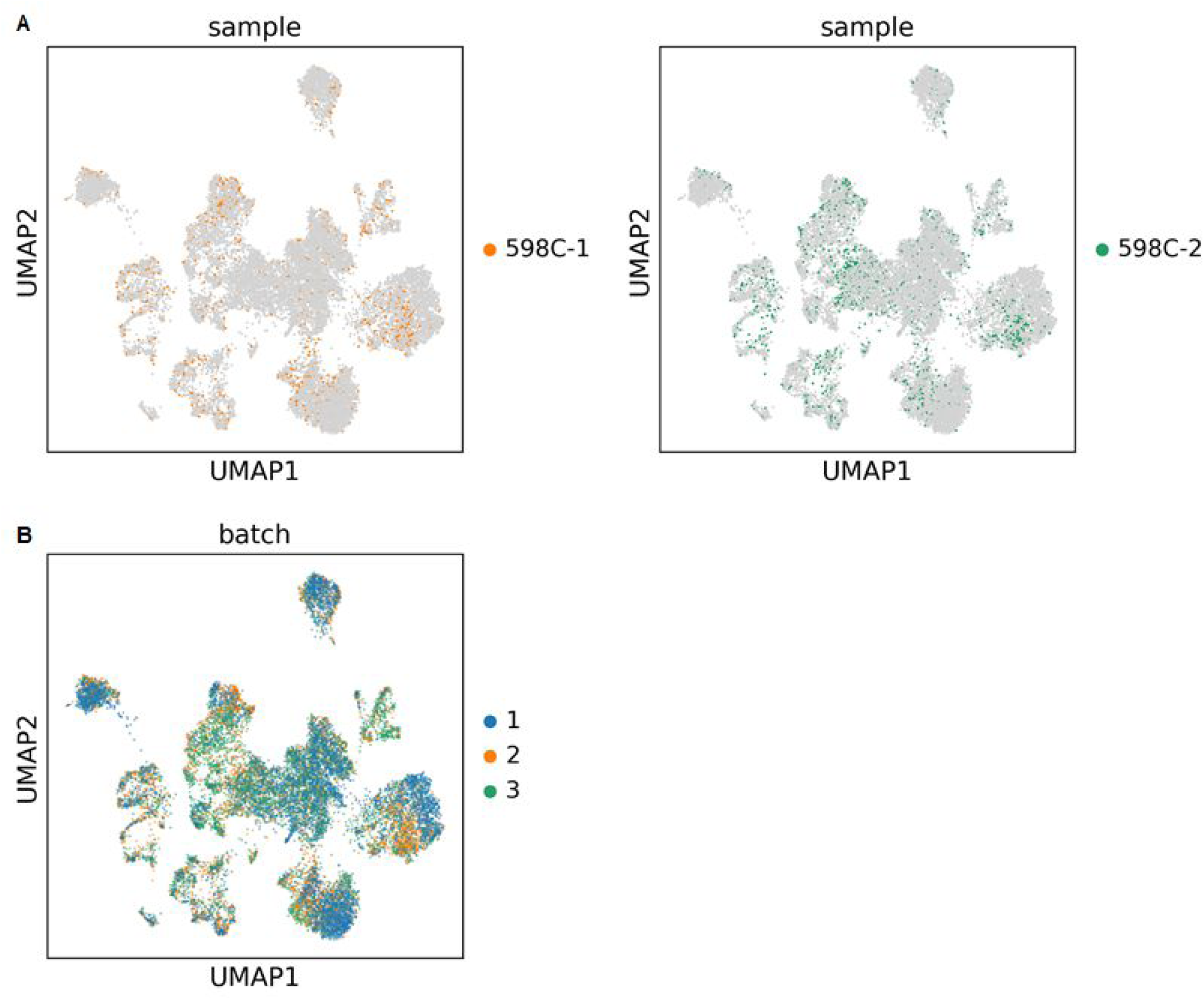
**(A)** UMAP plots highlighting nuclei from two technical replicate samples. Immediately after homogenizing the tissue from donor 598, the homogenate was split into two different aliquots (598C-1 and 598C-2). **(B)** UMAP plot highlighting nuclei based on the batch of samples from which they were isolated.

**Supplementary figure 2.**
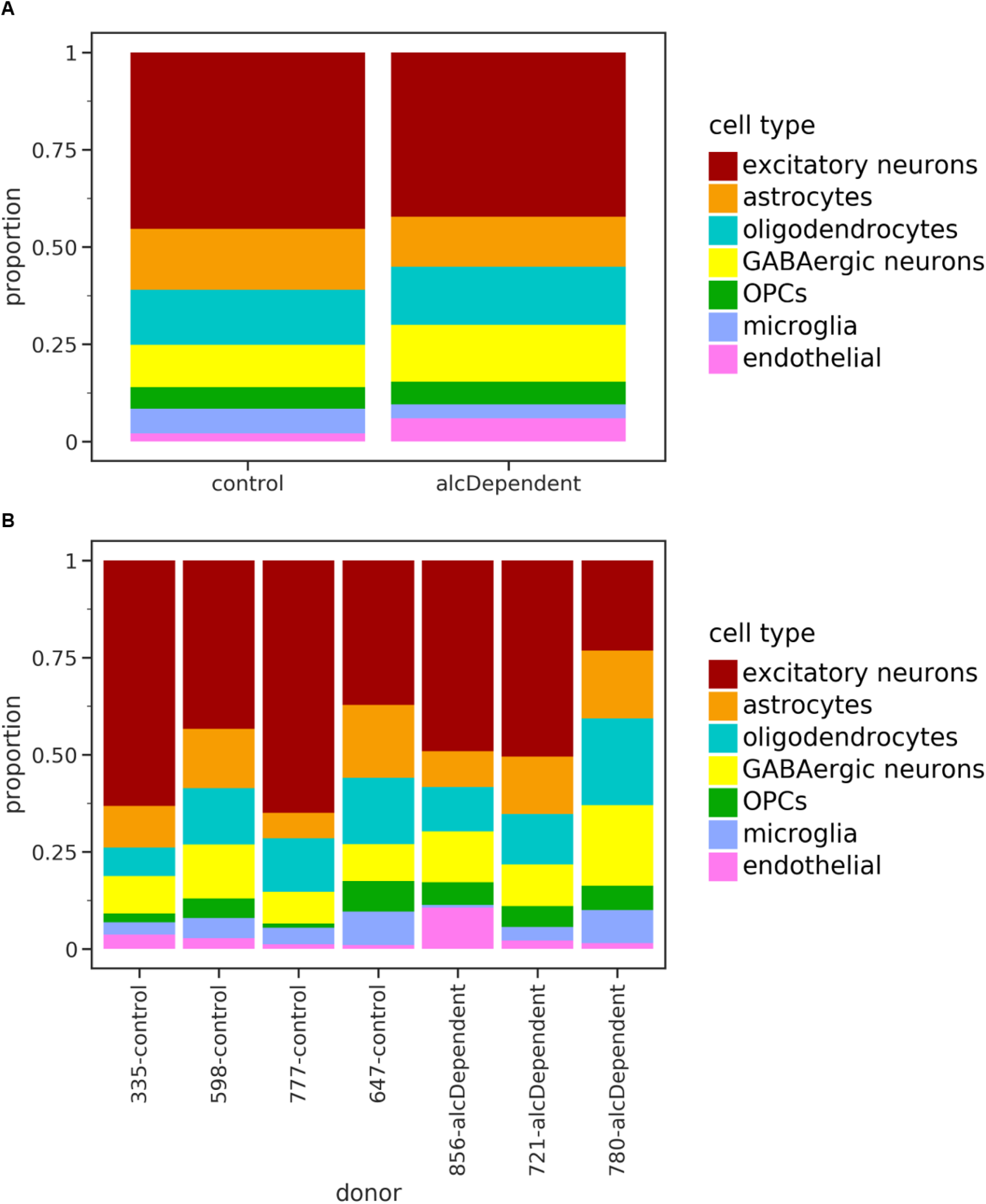
**(A)** Cell type proportions for nuclei from control and alcohol-dependent individuals. **(B)** Cell type proportions for nuclei from each specific donor. There were no significant differences in the proportion of any cell type between alcoholics and controls, based on t-tests. The lowest p-value was 0.27 for GABAergic neurons.

**Supplementary table 1.**
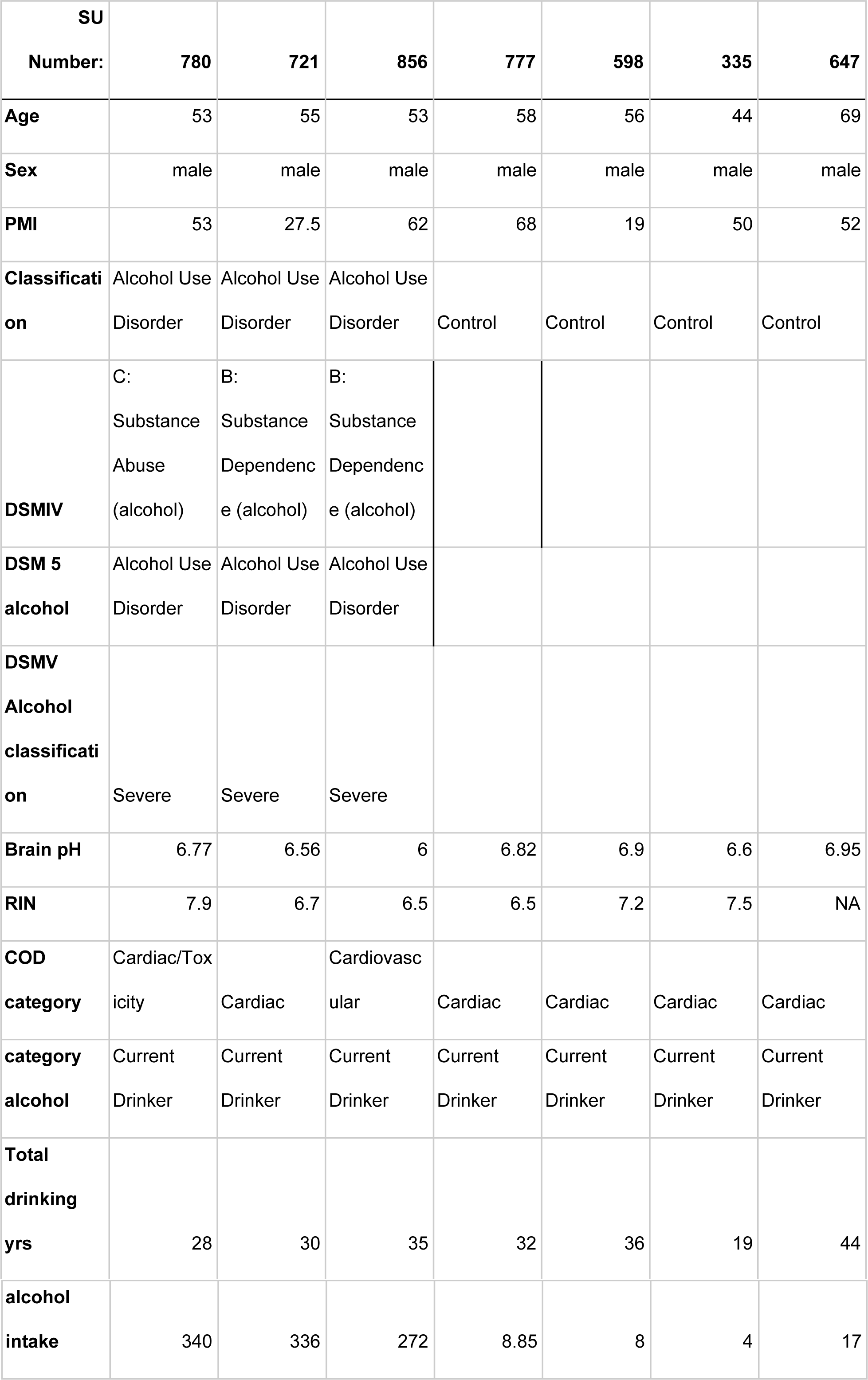
Donor information.

**Supplementary table 2.**
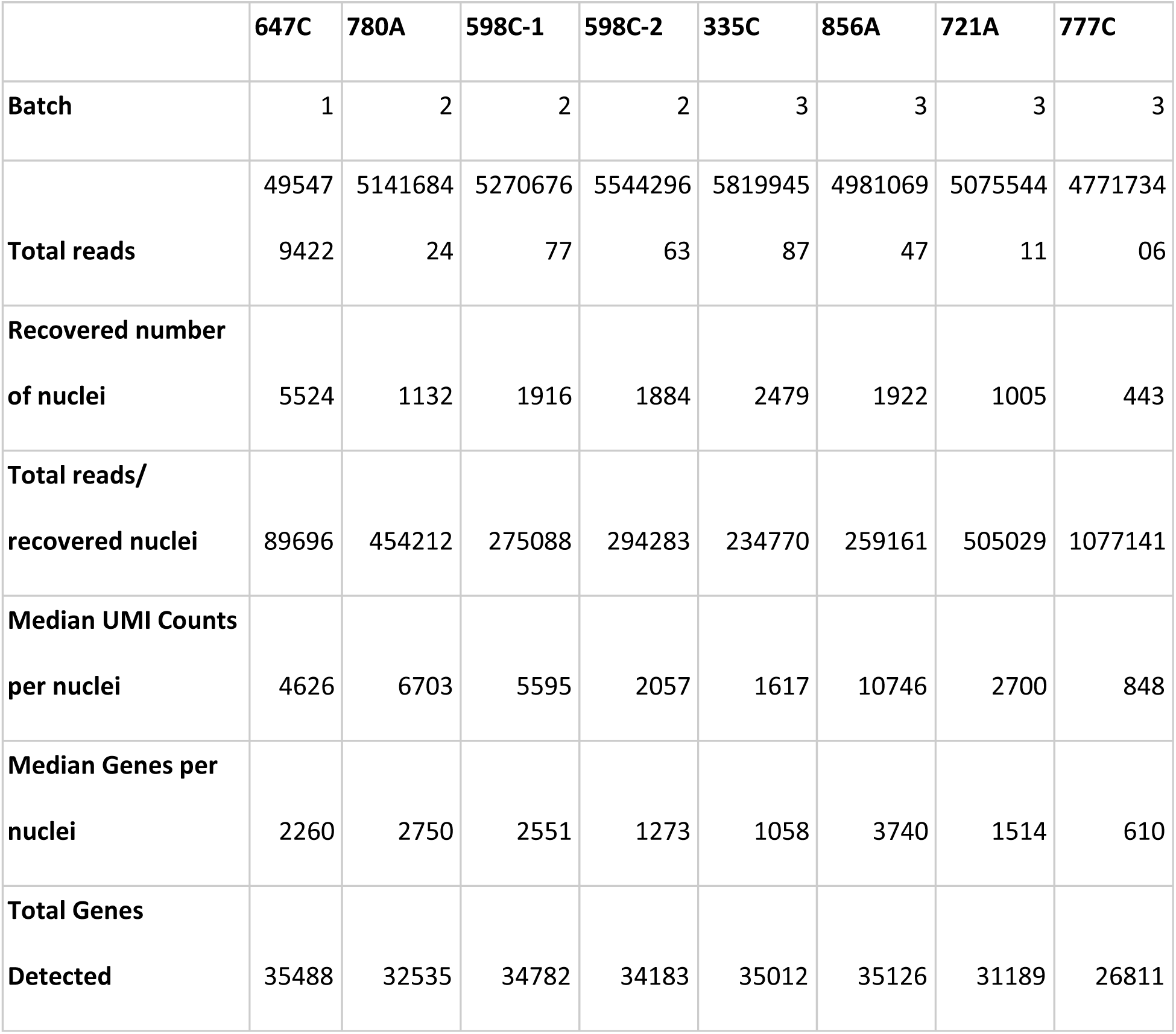
Technical information and quality control metrics for the snRNA-seq data.

